# Phylogenomics of montane frogs of the Brazilian Atlantic Forest supports a scenario of isolation in sky islands followed by relative climatic stability

**DOI:** 10.1101/226159

**Authors:** Marcio R. Pie, Brant C. Faircloth, Luiz F. Ribeiro, Marcos R. Bornschein, John E. McCormack

**Affiliations:** Departamento de Zoologia, Universidade Federal do Paraná, Curitiba, Paraná, Brazil; Mater Natura – Instituto de Estudos Ambientais, Curitiba, Paraná, Brazil; Department of Biological Sciences and Museum of Natural Science, Louisiana State University, Baton Rouge, Louisiana, USA; Escola de Ciências da Vida, Pontifícia Universidade Católica do Paraná, Curitiba, Paraná, Brazil; Instituto de Biociências, Universidade Estadual Paulista, São Vicente, São Paulo, Brazil; Moore Laboratory of Zoology, Occidental College, Los Angeles, California, USA

**Keywords:** *Melanophryniscus*, *Brachycephalus*, target enrichment, ultraconserved elements, coalescent

## Abstract

Despite encompassing a relatively small geographical area, montane regions harbor disproportionately high levels of species diversity and endemism. Yet, relatively little is known about the evolutionary mechanisms ultimately leading to montane diversity. In this study, we use target capture of ultraconserved elements to investigate the phylogenetic relationships and diversification patterns of *Melanophryniscus* (Bufonidae) and *Brachycephalus* (Brachycephalidae), two frog genera that occur in sky islands of the southern Atlantic Forest of Brazil. Specifically, we test whether diversification of montane species in these genera can be explained by a single climatic shift leading to isolation in sky islands, followed by relative climatic stability that maintained populations in allopatry. In both genera, the topologies inferred using concatenation and coalescent-based methods were concordant and had strong nodal support, except for a few recent splits. These recent splits tended to be supported by more informative loci (those with higher average bootstrap support), suggesting that, while individual trees may be well resolved, the relationships they recover are being obscured by non-informative data. Divergence dating of a combined dataset using both genera is consistent with concordant timing of their diversification. These results support the scenario of diversification-by-isolation in sky islands, and suggest that allopatry due to climatic gradients in montane regions are an important mechanism for generating species diversity and endemism in these regions.

## 1. Introduction

It has long been recognized that montane forests tend to display disproportionately high levels of species diversity and endemism, but relatively little is known about the processes underlying this phenomenon (Kessler and Kluge, 2008; Fjeldså et al., 2012). For example, diversification of montane lineages is often attributed to the direct effects of mountain uplift (e.g. Roy et al., 1997; Toussaint et al., 2014; Xing and Ree, 2017). However, the initial isolating mechanism may not simply result from the creation of a new, physical barrier. Rather, isolation could result from the intrinsic spatiotemporal heterogeneity in lineage persistence that subsequently produces high species turnover (Roy et al., 1997). Similarly, some researchers have argued that newly formed species arise in montane habitats and subsequently migrate into the lowlands, creating a situation where lowland habitats are ‘sinks’ of species accumulation rather than centers of species diversification (e.g. Fjeldså, 1994). For example, Roy et al. (1997) used distributional records of bird species in South America and Africa to suggest that avian diversification in these regions was driven by a dynamic process of local isolation in stable montane forests with occasional dispersal to other montane forest patches and that new species gradually expanded into other habitats, finally accumulating in the extensive tracts of lowland forest and woodland savannas. These examples demonstrate that we are still far from a comprehensive understanding of the relative contribution of different mechanisms generating and maintaining species diversity in montane regions and the importance of montane ecosystems as drivers of diversity across the landscape.

One unique model for investigating montane diversification is the Brazilian Atlantic Forest along the Serra do Mar mountain range. These mountains run parallel to the Atlantic coast of Brazil (Almeida and Carneiro, 1998) and form a barrier to moisture from the Atlantic (Safford, 1999a). This means that the South Atlantic anticyclone, a semi-permanent high pressure system that transports moist tropical air masses inland all year long (Behling, 2008), provides a constant source of precipitation, which is likely responsible for the formation of montane and cloud forests throughout the Serra do Mar mountain range (Behling, 2008). Along the Serra do Mar range, the interaction between geography and climate also produces dry environments caused by strong winds, with a thin soil layer and high levels of water-runoff, leading to the formation of high-elevation grasslands (*campos de altitude*) on many mountaintops (Safford, 1999a,b). These habitats are home to two anuran genera: *Melanophryniscus* (Bufonidae) and *Brachycephalus* (Brachycephalidae) (Figure 1).

**Figure 1.**
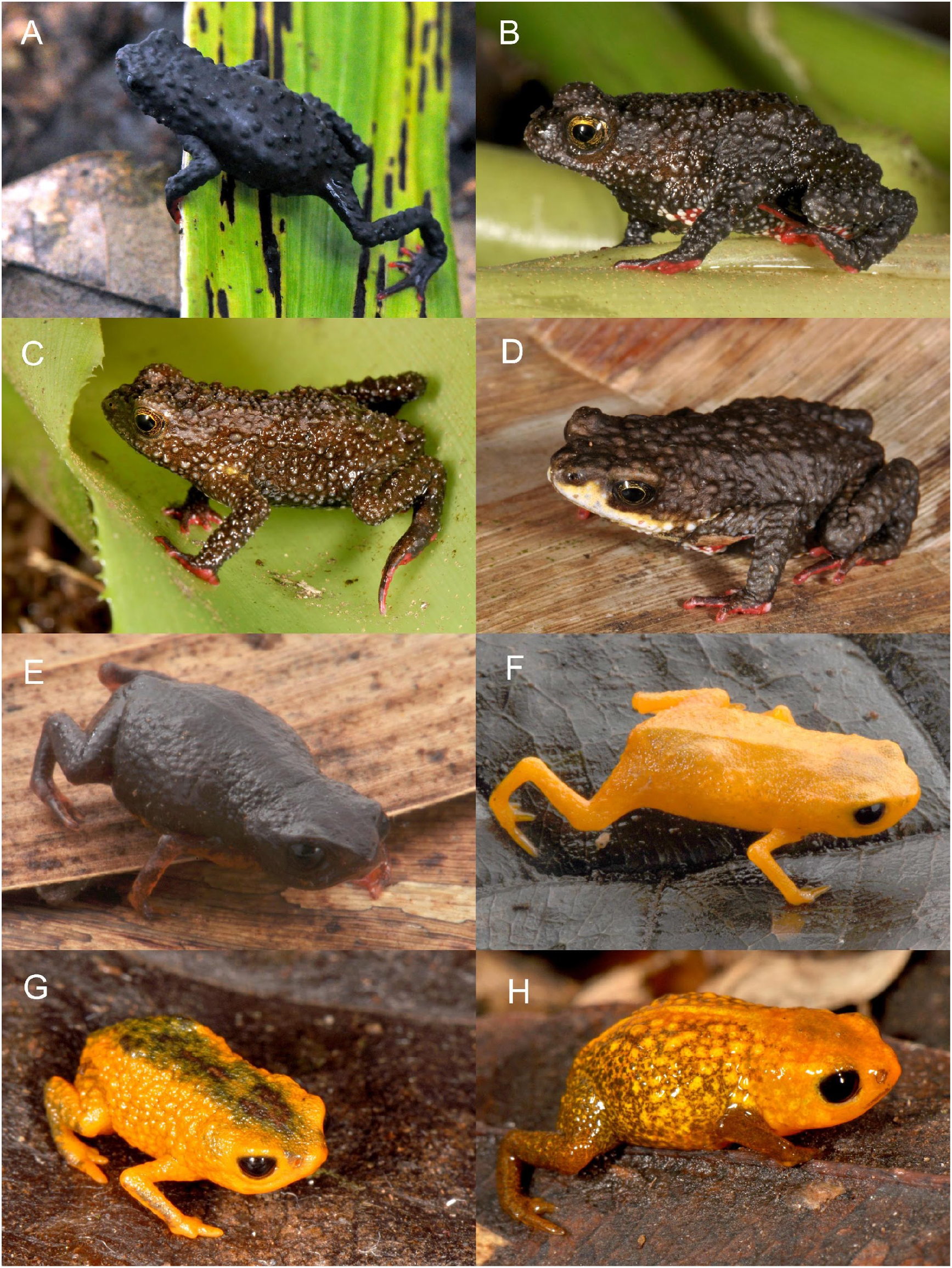
Examples of the species of *Melanophryniscus* and *Brachycephalus* investigated in the present study. A) *M. alipioi*; B) *Melanophryniscus* sp. Boa Vista; C) *M. milanoi;* D) *Melanophryniscus* sp. Morro do Boi; E) *B. brunneus;* F) *B. izecksohni;* G) *B. fuscolineatus;* H) *B. aurogutattus* (Photographs by LFR).

*Melanophryniscus* is widely distributed throughout southeastern South America, including parts of Brazil, Bolivia, Paraguay, Uruguay, and Argentina (Frost, 2017). The species of *Melanophryniscus* that occur in the Serra do Mar are montane endemics with restricted and isolated distributions in montane forests, cloud forests, *campos de altitude*, and inland grasslands (Langone et al., 2008; Steinbach-Padilha 2008; Bornschein et al., 2015), and include five of the 29 currently described *Melanophryniscus* species (Frost, 2017): *M. alipioi, M. biancae, M. milanoi, M. vilavelhensis*, and *M. xanthostomus*. These five species are unique among their congeners due to their reproduction in phytotelmata, as opposed to the other species that reproduce in temporary streamlets or temporary ponds (Baldo et al., 2014). Two of these species (*M. biancae* and *M. vilavelhensis*) represent a distinct lineage within montane *Melanophryniscus*, given their phylogenetic distinctiveness (Firkowski et al., 2016), their ecology (nocturnal versus diurnal), the unique type of vegetation in which they occur, and the plant species in which they reproduce (Bornschein et al., 2015). The actual number of the remaining montane *Melanophryniscus* is probably underestimated (Firkowski et al., 2016), because a recent phylogenomic species delimitation study suggests that some nominal species might be species complexes (Pie et al., 2017). *Brachycephalus* is endemic to the Brazilian Atlantic Forest, with a distribution extending nearly 1,700 km along the biome (Bornschein et al., 2016a). Most *Brachycephalus* species are found in isolated mountaintops, from the Brazilian states of Bahia in northeastern Brazil to Santa Catarina in southern Brazil (Pie et al., 2013; Pie and Ribeiro, 2015; Ribeiro et al., 2015; 2017; Bornschein et al., 2016a,b). The most remarkable feature of this genus is their extreme level of miniaturization (SVL≈1–1.5 cm), which has led to severe modifications of their body plan, such as the reduction in the number of digits (Hanken and Wake, 1993; Yeh, 2002). The distribution of montane species of *Melanophryniscus* and *Brachycephalus* overlap broadly across the southern Serra do Mar, with many pairs of species of each genus being found on the same mountains (Pie et al., 2013; Bornschein et al., 2015; 2016a; Firkowski et al, 2016). This condition of co-distributed microendemism provides a unique opportunity to investigate the timing of and the mechanisms driving their evolution. Concordant timing of diversification would be strong evidence for common vicariance processes.

A previous study of montane *Brachycephalus* and *Melanophryniscus* (Firkowski et al., 2016) used a small sample of loci to propose a two-step scenario explaining the diversification of montane species in the southern Serra do Mar. First, a climatic shift led cold-adapted species to track their ancestral niches (see Pie et al., 2013) into the mountaintops, creating a distributional arrangement commonly referred to as sky islands (McCormack et al., 2009). Then, climatic stability in the region maintained populations in isolation, leading to their diversification. The main sources of evidence for this scenario were twofold: the virtual lack of sympatry of microendemic species of each genus (Pie et al., 2013, Bornschein et al., 2015, 2016a) and the relative concordance in the timing of their diversification (Firkowski et al., 2016). However, the limited dataset used in Firkowski et al. (2016) left uncertainty in the estimated species trees upon which the inferred diversification scenario depended. Here, we use target capture of hundreds of ultraconserved elements (UCEs) to investigate the phylogenomic relationships and diversification patterns of *Melanophryniscus* and *Brachycephalus* in sky islands of the southern Serra do Mar in order to test whether they share a similar timing in their diversification, which would be consistent with common vicariance mechanisms leading to their isolation.

## 2. Material and methods

We obtained tissue samples from field-collected specimens of seven species of *Melanophryniscus* and 16 species of *Brachycephalus* (Table 1; see Pie et al. [2017] for a detailed account of the species delimitation, including still undescribed species). We deposited voucher specimens in the herpetological collection of the Department of Zoology of the Universidade Federal do Paraná (DZUP) and in the Museu de História Natural Capão da Imbuia (MHNCI), both in Curitiba, Brazil. More information on specimen collection methods and localities can be found in Firkowski et al. (2016). We extracted genomic DNA using a PureLink Genomic DNA kit (Invitrogen, USA), and we fragmented the obtained DNA using a BioRuptor NGS (Diagenode). We prepared Illumina libraries using KAPA library preparation kits (Kapa Biosystems) and ligated adapters to each sample that included unique, custom indexes (Faircloth and Glenn, 2012). To enrich targeted UCE loci, we followed an established workflow (Gnirke et al., 2009; Blumenstiel et al., 2010) while incorporating several modifications to the protocol detailed in Faircloth et al. (2012). Specifically, we pooled eight indexed libraries at equimolar ratios, prior to enrichment, we enriched each pool using a set of 2,560 custom-designed probes (MYcroarray, Inc.) targeting 2386 UCE loci (see Faircloth et al. [2012] and http://ultraconserved.org (last accessed August 5, 2014) for details on probe design), and we blocked the Illumina TruSeq adapter sequence using custom blocking oligos (Inosine at each index position in the blocking oligo). Prior to sequencing, we qPCR-quantified enriched pools, combined enriched pools at equimolar ratios, and sequenced the combined libraries using two, partial (50%) runs of a MiSeq PE250 (Cofactor Genomics). We performed species delimitation analyses using these data in a separate manuscript (Pie et al., 2017). Sequence reads for this project are available from NCBI BioProject PRJNA391191.

**Table 1.** Samples used in the present study. Unnamed species are indicated by “sp.”, followed by a code indicating the first recorded location.

| Species | Coordinates | Locality |
| --- | --- | --- |
| *Melanophryniscus alipioi* | 24°30'25"S, 48°48'49"W | Base of the Serra Água Limpa, municipality of Apiaí, São Paulo |
| *M. alipioi* | 25°07'49"S, 48°49'15"W | Capivari Grande, Serra do Capivari, municipality of Campina Grande do Sul, Paraná |
| *M. alipioi* | 24°51'17"S, 48°43'43"W | Caratuval, near the Parque Estadual das Lauráceas, municipality of Adrianópolis, Paraná |
| *M. alipioi* | 25°14'44"S, 48°50'04"W | Itapiroca, municipalities of Campina Grande do Sul, Paraná |
| *M. milanoi* | 26°47'55"S, 48°55'55"W | Morro do Baú, municipality of Ilhota, Santa Catarina |
| *M. milanoi* | 26°46'42"S, 49°01'57"W | Morro do Cachorro, on the border between the municipalities of Blumenau, Gaspar, and Luiz Alves, Santa Catarina |
| *M. xanthostomus* | 26°08'48"S, 49°10'43"W | Condomínio Vale dos Lagos, municipality of Joinville, Santa Catarina |
| *M. xanthostomus* | 26°01'17"S, 48°59'47"W | Serra do Quiriri, municipality of Campo Alegre, Santa Catarina |
| *Melanophryniscus* sp. “Azul” | 26°45'49"S, 49°12'23"W | Morro Azul, on the border between the municipalities of Pomerode and Rio dos Cedros, Santa Catarina |
| *Melanophryniscus* sp. “Boa Vista” | 26°30'58"S, 49°03'14"W | Morro Boa Vista, on the border between the municipalities of Jaraguá do Sul and Massaranduba, Santa Catarina |
| *Melanophryniscus* sp. “Boi” | 26°24'53"S, 49°13'08"W | Morro do Boi, municipality of Corupá, Santa Catarina |
| *Melanophryniscus* sp. “Igreja” | 25°36'40"S, 48°51'22"W | Morro dos Padres, Serra da Igreja, municipality of Morretes, Paraná |
| *Melanophryniscus* sp. “Igreja” | 25°37'31"S, 48°41'28"W | Torre da Prata, boundary of the municipalities of Morretes, Paranaguá, and Guaratuba, Paraná |
| *Brachycephalus auroguttatus* | 26°00'21"S, 48°55'25"W | Pedra da Tartaruga, municipality of Garuva, Santa Catarina |
| *B. boticario* | 26°46'42"S, 49°01'57"W | Morro do Cachorro, boundary of the municipalities of Blumenau, Gaspar, and Luiz Alves, Santa Catarina |
| *B. brunneus* | 25°13'29"S, 48°51'17"W | Abrigo 1, Serra dos Órgãos, municipality of Campina Grande do Sul, PR |
| *B. brunneus* | 25°15'59"S, 48°50'16"W | Camapuã, Serra dos Órgãos, boundary of the municipalities of Campina Grande do Sul and Antonina, Paraná |
| *B. brunneus* | 25°14'33"S, 48°50'04"W | Caratuva, Serra dos Órgãos, municipality of Campina Grande do Sul, Paraná |
| *B. curupira* | 25°42'07"S, 49°03'44"W | Serra do Salto, Malhada District, municipality of São José dos Pinhais, Paraná |
| *B. curupira* | 25°30'33"S, 48°58'58"W | Morro do Vigia, municipality of Piraquara, Paraná |
| *B. ferruginus* | 25°27'03"S, 48°54'59"W | Olimpo, Serra do Marumbi, municipality of Morretes, Paraná |
| *B. fuscolineatus* | 26°47'58"S, 48°55'47"W | Morro do Baú, municipality of Ilhota, Santa Catarina |
| *B. izecksohni* | 25°37'25"S, 48°41'31"W | Torre da Prata, Serra da Prata, boundary of the municipalities of Morretes, Paranaguá, and Guaratuba, Paraná |
| *B. mariaeterezae* | 26°06'51"S, 49°03'45"W | Reserva Particular do Patrimônio Natural Caetezal, top of the Serra Queimada, municipality of Joinville, Santa Catarina |
| *B. olivaceus* | 26°24'42"S, 49°12'59"W | Morro do Boi, municipality of Corupá, Santa Catarina |
| *B. pernix* | 25°23'19"S, 49°00'15"W | Anhangava, Serra da Baitaca, municipality of Quatro Barras, Paraná |
| *B. pombali* | 25°36'40"S, 48°51'22"W | Morro dos Padres, Serra da Igreja, municipality of Morretes, Paraná |
| *B. quiririensis* | 26°01'17"S, 48°59'47"W | Serra do Quiriri, municipality of Campo Alegre, Santa Catarina |
| *B. sulfuratus* | 25°20'17"S, 48°54'56"W | Corvo, municipality of Quatro Barras, Paraná |
| *B. verrucosus* | 26°12'44"S, 48°57'29"W | Morro da Tromba, municipality of Joinville, Santa Catarina |
| *B. sp. "Canasvieiras"* | 25°36'58"S, 48°46'59"W | Serra Canasvieiras, boundary of the municipalities of Guaratuba and Morretes, Paraná |
| *B. sp. "Tupipiá"* | 25°15'13"S, 48°48'20"W | Pico Tupipiá, Serra dos Órgãos, municipality of Antonina, Paraná |

We demultiplexed reads using automated procedures of the BaseSpace platform, and we filtered reads for adapter contamination, low-quality ends, and ambiguous bases using an automated pipeline (https://github.com/faircloth-lab/illumiprocessor) that incorporates Trimmomatic (Bolger et al., 2014). We assembled reads for each individual using Trinity (Grabherr et al., 2011). We used the PHYLUCE software package (Faircloth, 2015) to align assembled contigs back to their associated UCE loci, remove duplicate matches, create a taxon-specific database of contig-to-UCE matches, and extract UCE loci for all *Brachycephalus* and *Melanophryniscus* individuals. We also used PHYLUCE to harvest UCE loci from the *Xenopus* and *Rana* genomes for use as outgroup sequences. We then generated three sets of data for phylogenetic and divergence-time analyses: (1) all *Brachycephalus* species using *Melanophryniscus* sp. (collected in Morro dos Padres, Serra da Igreja, municipality of Morretes, Paraná) as the outgroup; (2) all *Melanophryniscus* species using *Brachycephalus sulfuratus* as the outgroup; and (3) a combined dataset including both *Brachycephalus* and *Melanophryniscus* species using *Xenopus* and *Rana* as outgroups. We filtered the loci included in each dataset to ensure there were no missing data, we aligned data for each individual in each data set using MAFFT (Katoh, 2013), and we trimmed resulting alignments using GBlocks (Castresana, 2000) with default parameters.

Following alignment, we carried out phylogenetic inference using both concatenated and coalescent-based methods. Concatenated analyses were carried out in RAxML 8.2.8 (Stamatakis, 2014) using a single GTRGAMMA model across the entirely of the concatenated data, and we performed 1000 rapid bootstrap replicates as a measure of branch support. We then inferred individual gene trees using the same parameters on RAxML and we performed coalescent-based phylogenetic inference using ASTRAL-II 5.0.3 (Mirarab and Warnow, 2015), a statistically consistent approach under the multi-species coalescent model and therefore capable of handling potential incomplete lineage sorting (Mirarab et al., 2014). Branch support was estimated using local posterior probabilities (LPPs, Erfan and Siavash [2016]).

We used RelTime (Tamura et al., 2012), as implemented in MEGA (Kumar et al., 2016) to estimate divergence times for the combined dataset, using a GTR+Γ5 model of evolution and a calibration between *Rana* and the ingroup at 147-162 Mya (Hedges et al., [2006], last accessed on April 1, 2017). RelTime first transforms an evolutionary tree with branch lengths, in the units of number of substitutions per site, into an ultrametric tree with relative times by estimating branch-specific relative rates for descendants of each internal node. This procedure is based on the fact that the time elapsed from the most recent common ancestor of two sister lineages is equal when all the taxa are contemporaneous (Tamura et al., 2012). RelTime then converts the ultrametric tree into a timetree using one or more calibration points (Kumar and Hedges, 2016). Comparisons between RelTime and a variety of large-scale datasets showed very high correlations in divergence time estimates when compared to alternative dating approaches such as MCMCTree and BEAST, but at a small fraction of the computation time (Mello et al., 2016). We then pruned the timetrees to include only one sequence per species in each genus, and we visualized the timing of lineage splits in each genus using lineage-through-time plots as implemented in the APE 4.1 (Paradis et al., 2004) package. We also estimated divergence times using BEAST v. 2.5.6 (Drummond et al., 2012, Bouckaert et al. 2014). We used an uncorrelated log-normal relaxed clock model (UCLN) with a calibrated Yule model for the tree prior and default priors for the remaining parameters and empirical nucleotide frequencies. To minimize computational demands, we used an unpartitioned GTR model and only one tip for each species. We used the same secondary calibration point indicated above, yet we emphasize that these estimates should be interpreted with caution, given that they can cause unrealistically narrow confidence intervals (Drummond & Bouckaert, 2014). We ran two replicates of each analysis for 100 million generations, sampling every 10,000 generations, and later combined separate runs using logcombiner v.2.4.7 (Bouckaert et al., 2014) and examined their combined log files in tracer v1.6 (Rambaut and Drummond, 2007) to assess convergence and burnin. All estimated parameters had ESS>400. Finally, the spatial distribution of the studied lineages was visualized with respect to their topography and divergence times using geophylogenies, as implemented in GenGIS 2.5.3 (Parks et al., 2013).

## 3. Results

After filtering loci with missing data, our final datasets included 820, 1,227, and 303 loci (total of 385,080, 677,667, and 155,683 bp) for the *Brachycephalus, Melanophryniscus*, and combined datasets, respectively, with 610 loci in common between the first two datasets. There was an increase in variability towards the flanking regions of the UCE core (Figure S1), with only 1.6 % and 13.2 % of the loci being invariant in the *Brachycephalus* and *Melanophryniscus* datasets, respectively (Figure S2). Basic properties of the UCE loci obtained in the present study are shown in Figure 2.

**Figure 2.**
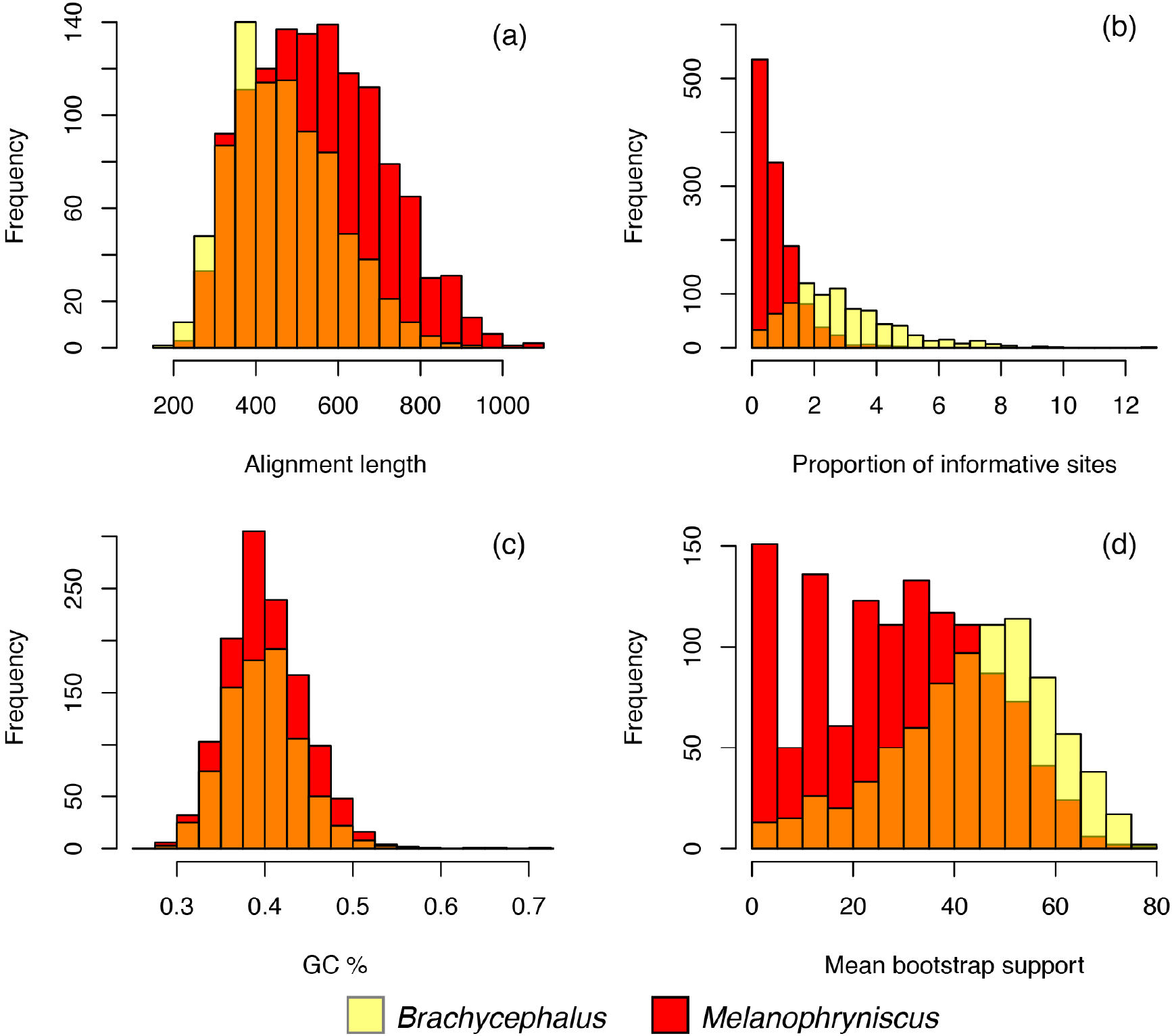
Properties of the ultraconserved element loci obtained for *Brachycephalus* and *Melanophryniscus*. Frequency distributions summarize the properties of the phylogenomic datasets on a per locus basis, including (a) alignment length; (b) proportion of parsimony-informative sites; (c) GC content; and (d) mean bootstrap support.

All phylogenetic analyses supported the same topologies (Figure 3), with slight differences in nodal support. Phylogenetic analyses of the concatenated *Brachycephalus* and *Melanophryniscus* datasets using ML provided 100 *%* bootstrap support for all nodes, except for the clade including *M. milanoi* and *M*. sp. “Azul”, which had a bootstrap support of 97 %. The concatenated ML analysis of the combined dataset provided the same topology as the previous datasets, but with decreased support at some of the nodes (Figure S3). Coalescent-based analyses also tended to show high support (LPP=0.94-1.0) for most nodes, except for three clades: *B. auroguttatus* + *B. quiririensis* (LPP=0.58), *B. verrucosus* + *B. olivaceus* (LPP=0.87), and *M. milanoi* + *M*. sp. “Azul” (LPP=0.88). To explore this issue further, we compared three statistics (average within-locus bootstrap support, proportion of informative sites, and fragment length) of the loci that were consistent with those clades in relation to the remaining loci (*B. auroguttatus* + *B. quiririensis:* supported by 126 loci, not supported by 694 loci; *B. verrucosus* + *B. olivaceus:* supported by 136 loci, not supported by 684 loci; *M. milanoi* + *M*. sp. “Azul”, supported by 595 loci, not supported by 225 loci) using Wilcoxon rank sum tests. In all three cases, the loci supporting those clades had significantly higher proportions of informative sites and mean bootstrap values (p=0.0015-6.11e-6) than the loci conflicting with these relationships, but not larger fragment lengths (p=0.53–0.053).

**Figure 3.**
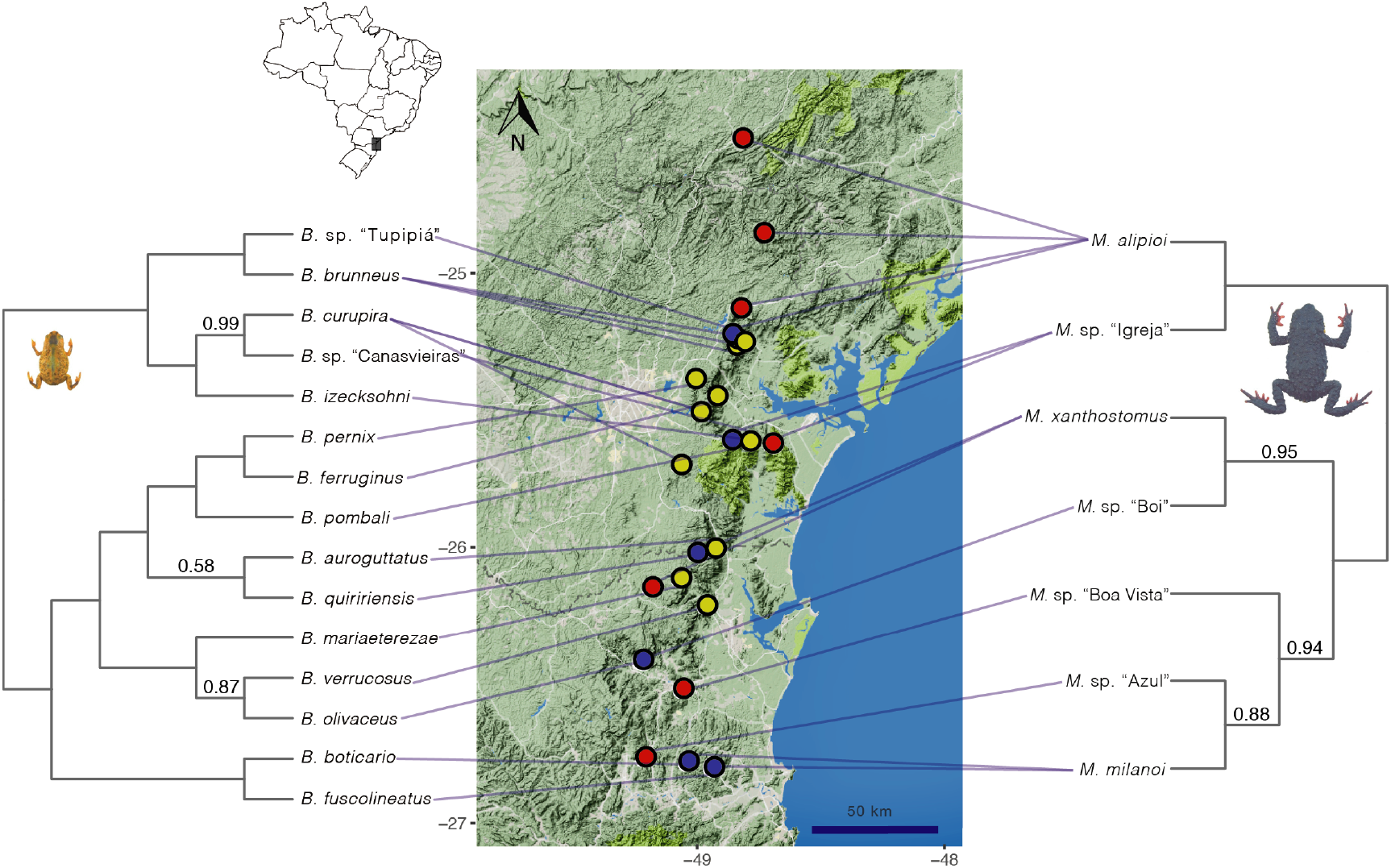
Relationships among the studied species of *Brachycephalus* (left) and *Melanophryniscus* (right). The topologies were identical in all concatenated and species tree analyses. Nodes above branches correspond to local posterior probabilities (LPPs) based on ASTRAL-II species tree analyses. Nodes without annotation are supported with LPP=1. All nodes were received 100% bootstrap in concatenated analyses (RAxML and BEAST). Outgroup lineages were omitted to facilitate visualization. Yellow and red circles indicate the presence of *Brachycephalus* and *Melanophryniscus*, whereas blue circles indicate locations where both genera were sampled.

RelTime and BEAST provided congruent estimates of divergence times (Figure 4, Figure S4), indicating that most species in both genera appear to have originated abruptly during the Pliocene (Figure 5). *Melanophryniscus* estimates were slightly older in RelTime in relation to BEAST, possibly because the latter were based on a matrix in which all but one tip per species was sampled prior to estimation. The concordant timing of the recent divergence times in each genus were consistent with a scenario in which their speciation events were driven by the same isolation mechanisms (Figure 4). On the other hand, despite the strong temporal congruence in the diversification of both genera, they did not share a common geographical distribution. For instance, *Melanophryniscus* species were distributed across a simple north-south axis, whereas there was some overlap between the distributions of the two major clades of *Brachycephalus* in the state of Paraná (Figure 3). Interestingly, this overlap is associated with some of the most recent speciation events in *Brachycephalus* (Figure 3).

**Figure 4.**
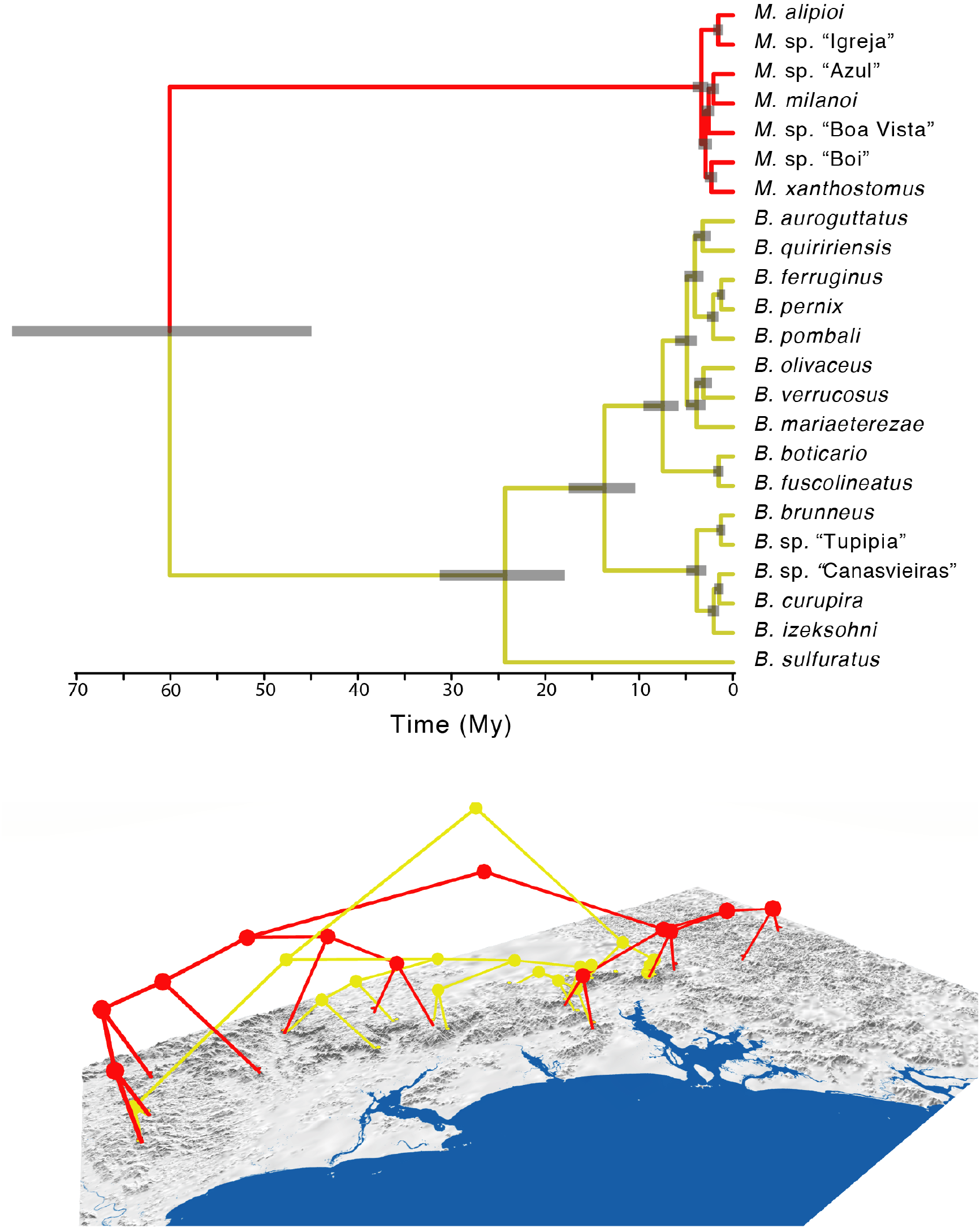
Timing of diversification of *Brachycephalus* (yellow) and *Melanophryniscus* (red) and their distribution over geographical space. Divergence time estimates are based on 150 million MCMC generations of an unpartitioned matrix (303 UCE loci, 155,683 bp) under a GTR+Γ4 model of evolution. Divergence estimates based on RelTime are indicated in Figure S4. The distribution of the studied lineages and their relationships are shown in relation to the topography of the region (below; see Figure 3 for scale).

**Figure 5.**
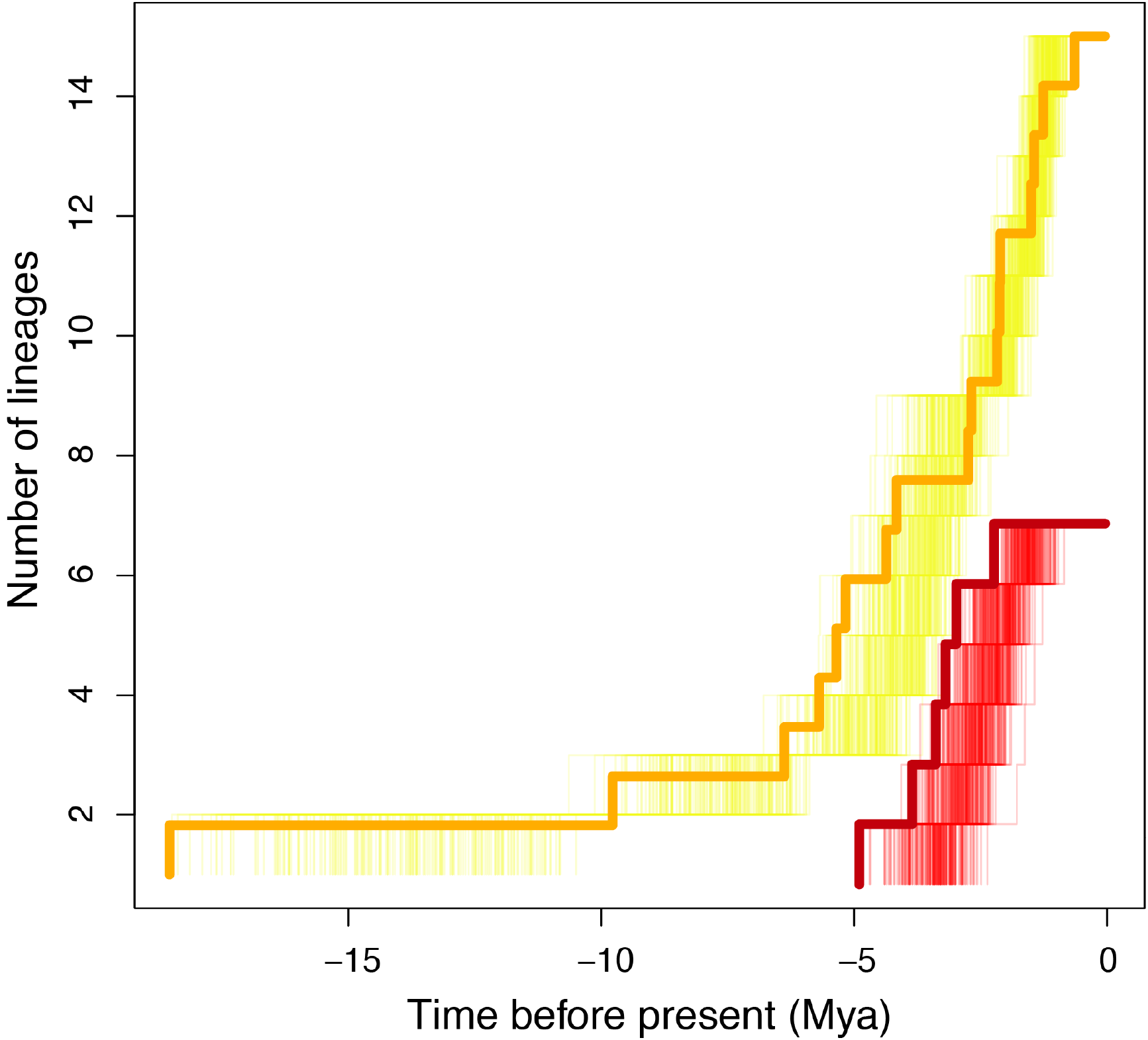
Lineages-through-time plots indicating the timing of diversification of *Brachycephalus* (yellow) and *Melanophryniscus* (red). Thin lines correspond to 200 post-burnin trees from BEAST analyses, whereas thick lines indicate divergence times based on the RelTime method.

## 4. Discussion

We observed substantial differences in the phylogeny inferred using UCE data compared to that of Firkowski et al. (2016). *Melanophryniscus xanthostomus* and *M*. sp. “Boi” were shown to be more closely related to the clade including *M. milanoi* and related lineages than to *M. alipioi* and *M*. sp. “Igreja”. In addition, we detected many differences in the southern clade of *Brachycephalus*, particularly with respect to the phylogenetic position of *B. auroguttatus* and *B. quiririensis*. These differences may have resulted from the rapid diversification of these lineages (Figure 4), which can require large-scale datasets to be resolved (Smith et al., 2015). The consistency in the obtained topologies across methods indicates that our results are likely to be a solid basis for further interpretation of the diversification patterns in these anurans.

Estimates of divergence times in this study are older than those from a previous study that used anuran ND2 mutation rates (Firkowski et al., 2016). Indeed, our estimates bring the speciation events in the studied *Brachycephalus* and *Melanophryniscus* closer to divergences found in other montane lineages (e.g. Toussaint et al., 2014). On the other hand, geological processes that gave rise to the Serra do Mar predates these speciation events considerably, given that the Serra do Mar was formed by the differential erosion of rocks of varying levels of resistance that took place from Paleogene through the Miocene (Almeida and Carneiro, 1998). The inference of older divergence times for both lineages creates another conundrum: how was it possible for these species to have retained microendemism and allopatry, given habitat change and potential connectivity of mountains over that time? It seems unlikely that each lineage remained isolated without either dispersing or colonizing nearby areas over this long period of time, especially given that some are less than a few km distant from one another.

According to the scenario proposed in Firkowski et al. (2016), this could have been achieved through climatic stability, which prevented colder climates from expanding into lowlands, thus hampering the possibility of secondary contact and sympatry (see Wiens [2004 and Kozak and Wiens [2006; 2010] for a general discussion of how niche conservatism might lead to population isolation and divergence). While stability in montane habitats in this region might seem unlikely, given that grasslands covered many areas of the Atlantic Forest during glacial times (e.g. Behling, 2008), areas of paleoecological stability may have persisted in small pockets within mountains (Roy et al., 1997), thus acting as micro-refugia (sensu Brown and Ab’Saber, 1979).

The new phylogenies present here immediately reveal some fascinating evolutionary scenarios that could be followed up on by detailed study. For instance, most of the species in the northern clade of *Brachycephalus* (Figure 4a,c) are characterized by highly cryptic coloration, including a dark brown dorsum (e.g. *B. brunneus*, Figure 1E). However, one species (*B. izecksohni*, Figure 1F), and many other *Brachycephalus*, are highly aposematic, with bright coloration patterns warning of the presence of a powerful neurotoxin (e.g. Pires et al., 2002). The phylogenetic distribution of coloration patterns suggests that aposematism was lost at the origin of this clade, and later regained with the evolution of *B. izecksohni* (Ribeiro et al., 2017). It is also noteworthy that there are exceptions to the rule that most of the species of *Brachycephalus* in this study are only found in one or a few adjacent mountaintops - namely *B. brunneus* and *B*. *curupira*, which are precisely the species with cryptic coloration. Even broader geographical distributions are found in more distantly-related lineages in the *didactylus* group (*sensu* Ribeiro et al., 2015), such as *B. didactylus, B. hermogenesi* and *B. sulfuratus*, which are all cryptic. The wider distribution found in all cryptic *Brachycephalus* lineages might indicate that skin coloration patterns could play a role as one of the factors driving endemism in the genus.

## 5. Acknowledgments

This study was partially funded by a grant from Fundação Grupo Boticário de Proteção à Natureza (grant #A0010_2014). Samples were collected under ICMBio permit #22470-2 and Instituto Ambiental do Paraná permit 355/11. MRP was funded by CNPq/MCT (grant 301636/2016-8). Whitney Tsai provided invaluable assistance for obtaining UCE data.

---

**Figure S1.**
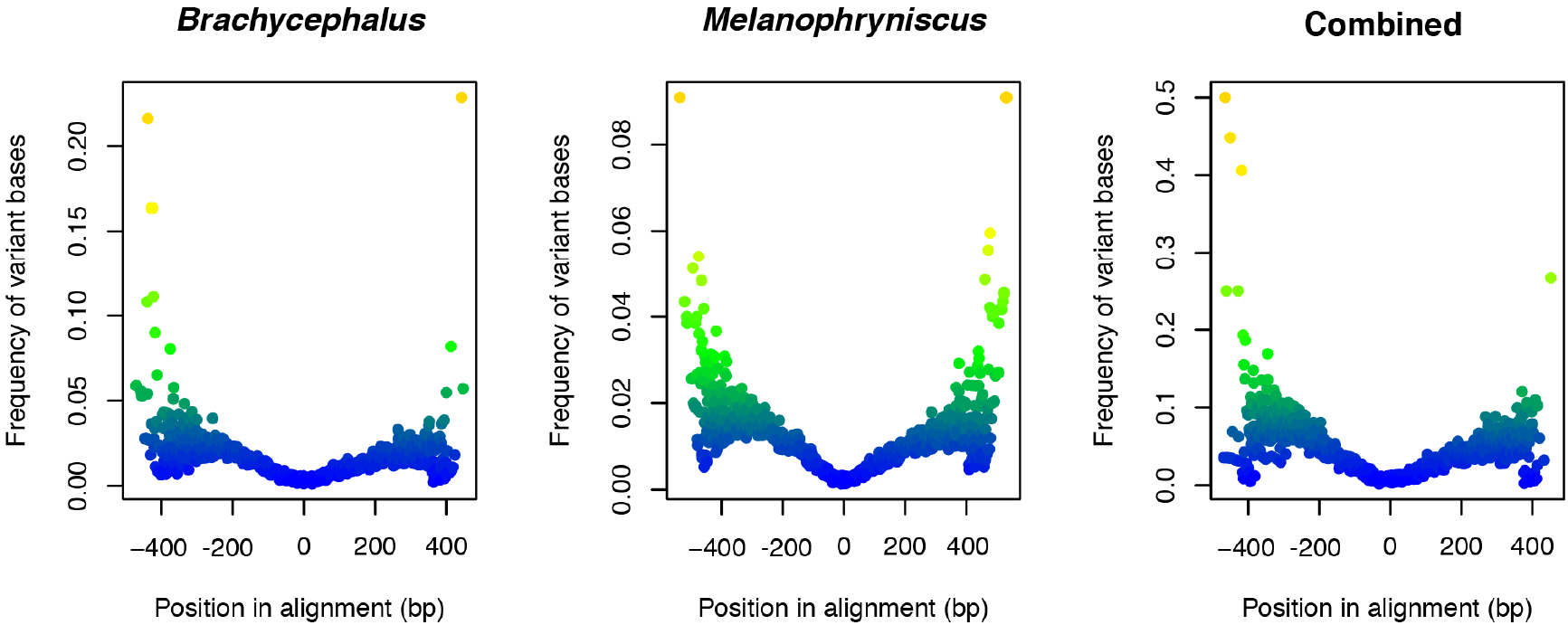
Increase in variability flanking the ultraconserved regions of the studied datasets.

**Figure S2.**
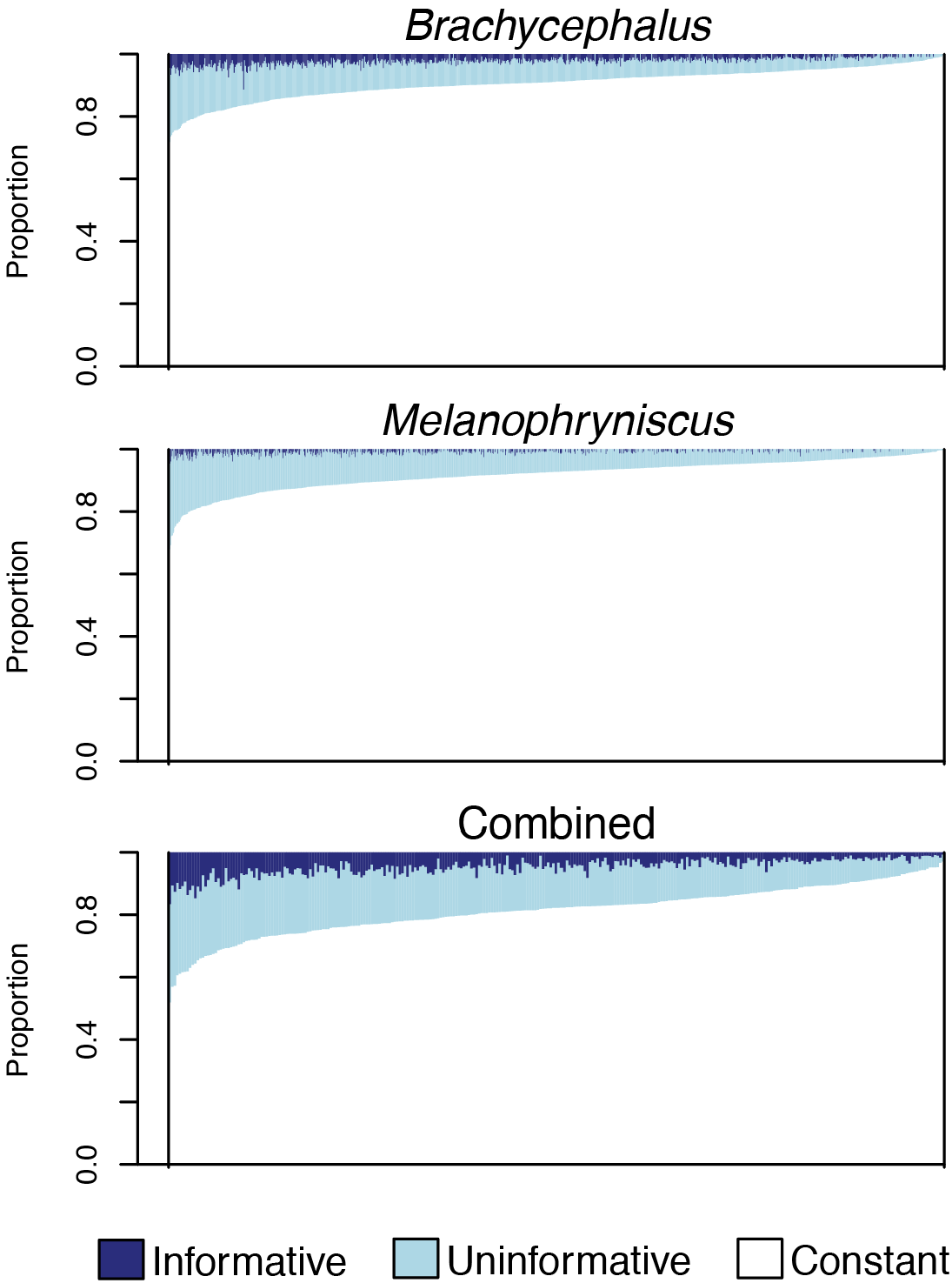
Proportion of parsimony-informative, parsimony-uninformative, and constant sites per locus in each dataset. Loci were ranked by the number of variable sites to facilitate visualization.

**Figure S3.**
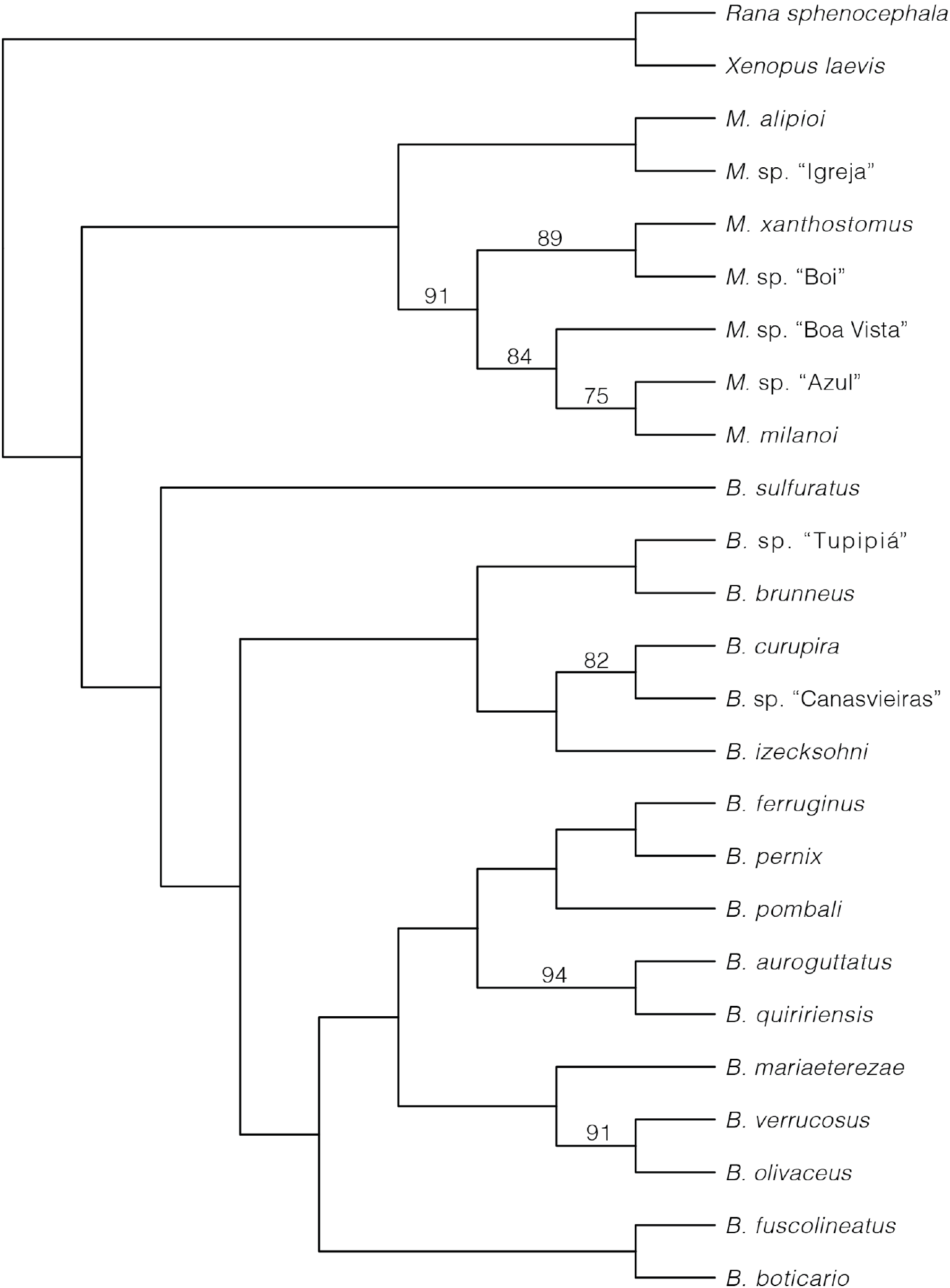
Phylogenetic relationships among the studied species, as inferred by the concatenated maximum likelihood analysis of the combined dataset. Values on branches indicate bootstrap support values (based on 1000 replicates). Branches without values are supported by 100% bootstrap support.

**Figure S4.**
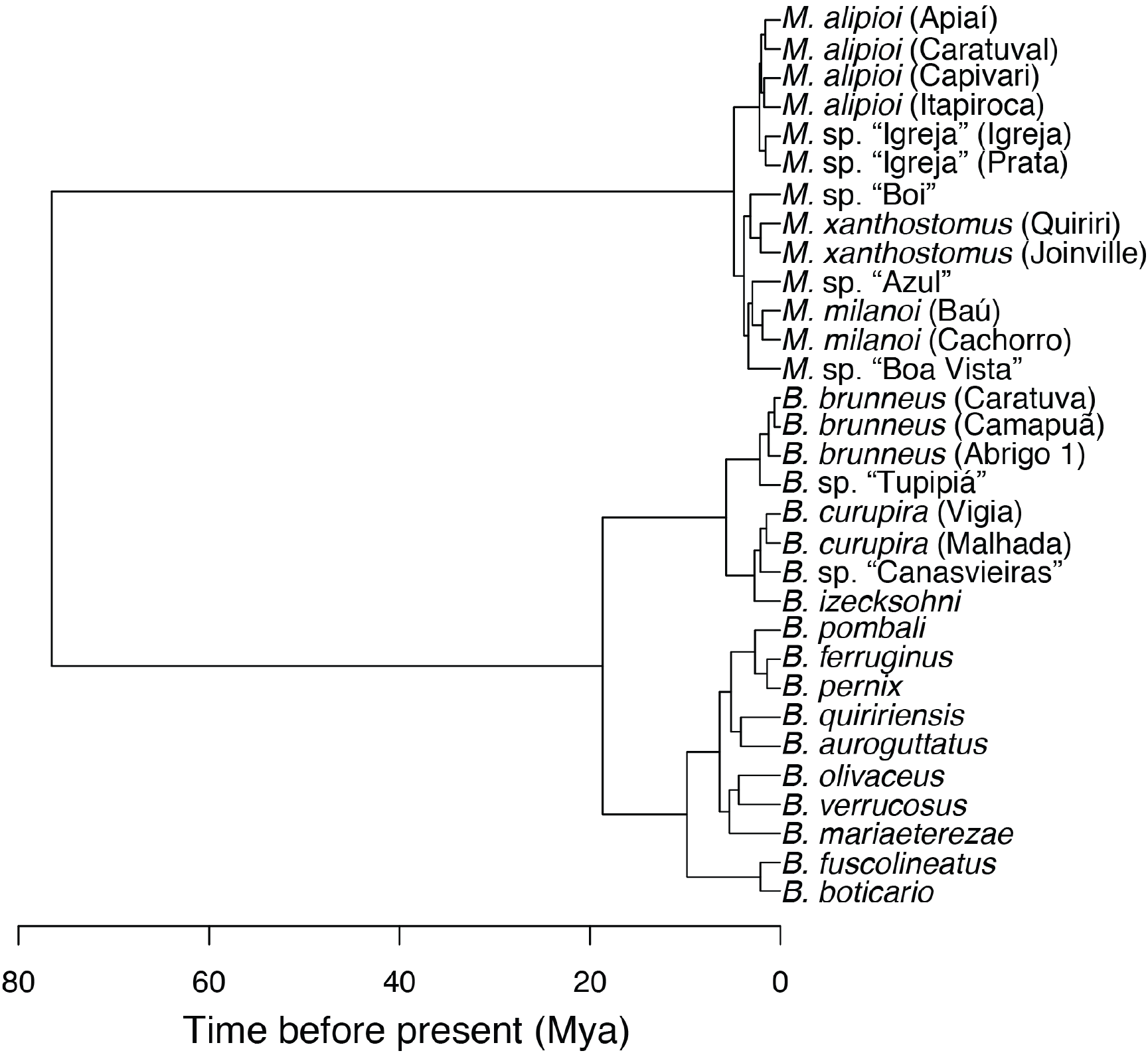
Phylogenetic relationships between studied species based on RelTime estimates of concatenated sequences of 303 UCE loci (155,683 bp).

